# Amplicon sequencing of variable 16S rRNA and ITS2 regions reveal honeybee susceptibility to diseases resulted of their dietary preferences under anthropogenic landforms

**DOI:** 10.1101/2021.01.28.428626

**Authors:** Aneta A. Ptaszyńska, Przemysław Latoch, Paul J. Hurd, Andrew Polaszek, Joanna Michalska-Madej, Łukasz Grochowalski, Dominik Strapagiel, Sebastian Gnat, Daniel Załuski, Marek Gancarz, Robert Rusinek, Patcharin Krutmuang, Raquel Martín Hernández, Mariano Higes Pascual, Agata L. Starosta

**Affiliations:** Department of Immunobiology, Institute of Biological Sciences, Faculty of Biology and Biotechnology, Maria Curie-Skłodowska University, Akademicka 19 Str., 20-033 Lublin, Poland; School of Biological and Chemical Sciences, Queen Mary University of London, London, E1 4NS, United Kingdom; Polish-Japanese Academy of Information Technology, Koszykowa 86 st., 02-008 Warsaw, Poland; Laboratory of Gene Expression, ECOTECH-Complex, Maria Curie-Sklodowska University, ul. Gleboka 39, 20-612 Lublin, Poland; Department of Life Sciences, Insects Division, Natural History Museum, London SW7 5BD UK; Biobank Lab, Department of Molecular Biophysics, Faculty of Biology and Environmental Protection, University of Łódź, Pilarskiego 14/16, 90-231 Łódź, Poland; University of Life Sciences, Faculty of Veterinary Medicine, Institute of Preclinical Veterinary Sciences, Department of Veterinary Microbiology, Akademicka 12, 20-033 Lublin, Poland; Department of Pharmaceutical Botany and Pharmacognosy, Ludwik Rydygier Collegium Medicum, Nicolaus Copernicus University, Marie Curie-Skłodowska 9, 85-094 Bydgoszcz, Poland; Institute of Agrophysics, Polish Academy of Sciences, Doświadczalna 4 Str., 20-290 Lublin, Poland; Faculty of Production and Power Engineering, University of Agriculture in Kraków, Balicka 116B, 30-149 Kraków, Poland; Department of Entomology and Plant Pathology, Faculty of Agriculture, Chiang Mai University, 50200, Thailand; Research Center of Microbial Diversity and Sustainable Utilization, Faculty of Science, Chiang Mai University, Chiang Mai 50200, Thailand; IRIAF. Instituto Regional de Investigación y Desarrollo Agroalimentario y Forestal, Laboratorio de Patología Apícola, Centro de Investigación Apícola y Agroambiental (CIAPA), Consejería de Agricultura de la Junta de Comunidades de Castilla-La Mancha, Camino de San Martín s/n, 19180 Marchamalo, Spain; Instituto de Recursos Humanos para la Ciencia y la Tecnología (INCRECYT-FEDER), Fundación Parque Científico y Tecnológico de Castilla—La Mancha, 02006 Albacete, Spain; Department of Molecular Biology, Institute of Biological Sciences, Maria Curie-Sklodowska University, Akademicka 19 Str., 20-033 Lublin, Poland

**Keywords:** *Apis mellifera*, 16S rRNA, ITR2, NGS, *Nosema apis*, *Nosema ceranae*, *Nosema bombi*, *Acarapis woodi*, Trypanosomatida, *Crithidia* spp., neogregarines, *Apicystis* spp., antropocene, insectageddon, urban area, urban environment, bee biology

## Abstract

European *Apis mellifera* and Asian *Apis cerana* honeybees, are essential crop pollinators. Microbiome studies can provide complex information on health and fitness of these insects in relation to environmental changes, and plant availability. Amplicon sequencing of variable regions of 16S rRNA and internally transcribed spacers (ITSs) allow identification of the metabiome. These methods provide a tool for monitoring otherwise uncultured microbes isolated from the gut of the honeybees. They also help monitor the composition of the gut fungi and, intriguingly, pollens collected by the insect. Here, we present data from amplicon sequencing of the 16S rRNA and ITS2 regions from honeybees collected at various time points from anthropogenic landforms as urban areas in Poland, UK, Spain, Greece, and Thailand. We have analysed microbial composition of honeybee intestine as well as fungi and pollens. We conclude that differences between samples were mainly influenced by the bacteria, plant pollens and fungi, respectively. Moreover, honeybees feeding on a honeydew diet, mainly based on sugars, were more prone to fungal pathogens (*Nosema ceranae*) and neogregarines. Finally, the period when honeybees switch to the winter generation (longer-lived forager honeybees) is the most sensitive to diet perturbations and hence pathogens attack, for the whole beekeeping season. It is possible that evolutionary adaptation of bees fails to benefit them in the modern anthropomorphised environment.

## 1. Introduction

The next-generation sequencing (NGS) is a culture-independent method often used for studding entire microbial communities and helps to understand how microbes influence health and diseases of humans and animals including a honeybee [1–3]. Adult honeybees harbor a specialized gut microbiota of relatively low complexity with diet as a major factor to differences in bacterial loads [4]. Although honeybee microbiome core species is quite consistent regardless of environmental, geographical and genetic differences between specimens [2], some studies indicated that it can be sensitive to infections, changes in diet, malnutrition and many anthropogenic activities, as extensive pesticide use and urban land-use changes [5–7].

Currently, in developed countries, anthropogenic landforms are the most impacting features and include landforms created either directly by human activity, or indirectly by natural processes triggered by human activity [8–10]. Not only has human activity influenced geological features, but it has also considerably affected flora and fauna [11]. The loss in biodiversity is often described as the sixth mass extinction and a slump in insects’ mass and biodiversity is so spectacular that the term Insectaggedon is appropriate to describe the trend [12–16].

Recent research shows insects to be dying out eight times faster than mammals, birds, or reptiles [12, 17, 18]. Most noteworthy factors behind the decline of insects are inappropriate application of pesticides, increased use of fertilizers and intense agronomic activities, highly intensive farming, insects’ malnutrition caused by farmland monocultures, parasites, long-term drought, long-term lack of sun, especially accompanied by low temperatures, as well as viral, bacterial, and fungal diseases [10, 19]. Nowadays, special concern is laid to declining of pollinators for their valuable ecosystem services [20]. Therefore, both for educational purposes and as a way to preserve pollinator populations is to keep bees in urban areas (i.e. programmes entitled: “Life+ Urbanbees” [21], “City Bees” [22], “Urban Beekeeping” [23]). The aim of the study was to use the amplicon sequencing of variable 16S rRNA and ITS2 to screen of honeybee colonies originated from different urban areas and to check if really “the bees love the city”.

## 2. Results and Discussion

Adult honeybees harbour specialized gut microbiota of relatively low complexity with five core bacterial strains [3, 4, 24, 25, 26]: *Lactobacillus* Firm-4 and Firm-5 (Firmicutes)*, Giliamella* (γ-proteobacteria, Orbales)*, Snodgrassella* (γ-proteobacteria, Neisseriaceae)*, Bifidobacterium* (Actinobacteria) and a number of elective bacterial strains, including *Frischella* (γ-proteobacteria, Orbales), *Bartonella* (α-proteobacteria, Rhizobiales), *Commensalibacter* (α-proteobacteria, Acetobacterales) and *Bombella* (α-proteobacteria, Acetobacterales), which was also confirmed in this study (Figure 1, Figure 2 1, Table 1, Table 2).

**Figure 1.**
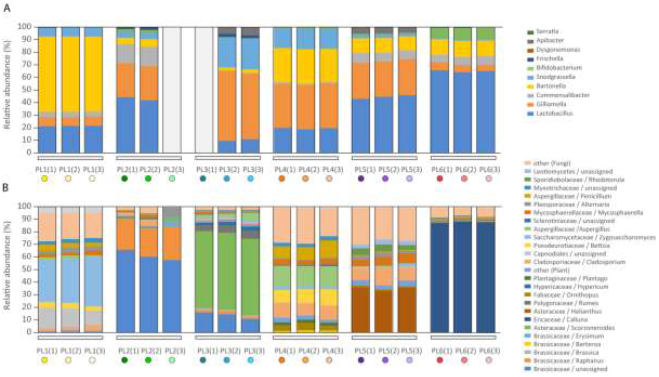
Composition of bacteria (A), fungi and pollen (B) form Polish honeybee samples.

**Figure 2.**
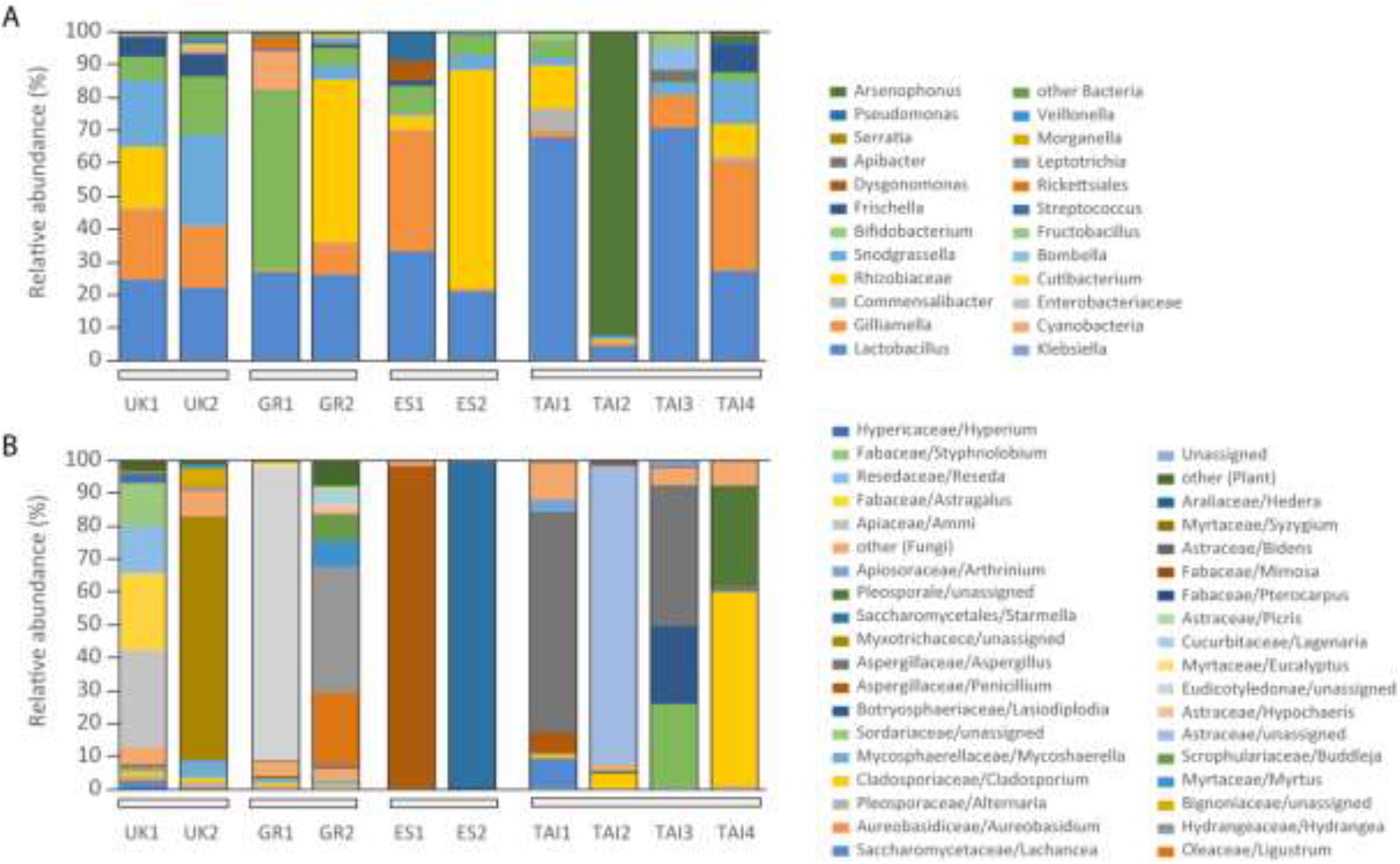
Composition of bacteria (A), fungi and pollen (B) form UK, Greek, Spanish, and Thai bee samples.

NGS was successfully used for taxonomic assessment of pollens and plants from many ecological and polynological studies, and to determine plant–pollinator interactions or to confirm the floral composition of honey [3, 27–30].

### 2.1. Microbiome and pollen composition of honeybees from Poland (differences over the vegetation season)

Following NGS analysis, it was possible to assign samples to the time that they were collected by comparing them with vegetation periods of nectarious and pollen-rich plants. From one location, 3 specimens (forager honeybees) were taken, as the representative and consistent number for each group (data adequacy confirmed by the PCA analysis of 16S and ITS2 amplicons, Supplementary materials Figure S1). Microbiome down to genera analysis helped to divide Polish honeybee samples into 6 sub-groups: PL1, PL2, PL3, PL4, PL5, and PL6 (Table 1).

Forager honeybees from PL1 group were collected in April and 16 taxa (species level) were identified in total from the analysed 16S amplicon and 164 taxa (species level) in ITS2 analyses (Supplementary materials Table S1). PL1 honeybees most likely foraged on honeydew, which was typified by the low number of pollens (1.81%, SD=0.512 belonging primarily to the *Betula* sp. and *Urtica* sp. genera) and a high number of fungal spores (93.83%, SD=0.554). A sugar-based diet, such as honeydew being the staple food, can led to higher yeast numbers as observed in PL1 (35.92%, SD=3.208) (Tab. 1 and 2). Furthermore, a pathogenic fungus belonging to *Nosema ceranae* was detected in the PL1 (Supplementary materials Table S2).

PL2 forager honeybees were collected in May and 16 taxa were identified in total from 16S amplicons analysed and 147 taxa in ITS2 (Supplementary materials Table S1). *Lactobacilli* genera were the dominant bacterial species in PL2 (43.02%, SD=1.704) (Tab. 1). At the same time, the load of fungal spores was moderate (9.59%; SD=4.476) with the prevalence of *Cladosporium* and a small content of spores from other genera (Tab. 2). The dominant pollen content in the PL2 with 87.54% (SD=3.886) came from plants from the *Brassicaceae* family, mainly unassigned and from the *Raphanus* genus (Tab. 1B) known for its high protein pollen content [31]. No pathogens were detected in PL2 (Supplementary materials Table S2).

PL3 forager honeybees were collected in June and 18 taxa were identified in total from 16S amplicons analysed and 177 taxa in ITS2 (Supplementary materials Table S1). This group shows a higher number of *γ*-proteobacteria from *Gilliamella* and *Snodgrassella* genera (54.03%, SD=2.333) (Tab. 1). PL3 honeybees collected pollen from Polygonaceae plants, which contain moderate amounts of amino acids [30]. In this group, fungal spore load was moderate (11.25%, SD=1.847) with the *Aspergillus, Cladosporium* and *Mycosphaerella* (mainly transferred as bioaerosols by wind in the air) at the level of 3.76% (SD=1.9486), 1.21% (SD=0.0961), and 0.81% (SD=0.624), respectively (Tab. 1B). No pathogens were detected in PL3 (Supplementary materials Table S2).

PL4 forager honeybees were collected in July and 16 taxa were identified in total from 16S amplicons analysed and 322 taxa in ITS2 (Supplementary materials Table S1). This group was differentiated on the basis of the highest fungal DNA loads (87.31%, SD=1.680) and trace amounts of plant DNA (Tab. 1B). The fungi derived mainly from the *Aspergillus* (15.98%, SD=0.503), *Cladosporium* (10.91%, SD=1.048), *Penicillum* (11.07%, SD=2.190), and *Betsia* (11.20%, SD=1.572) genera. The spores of the three first genera commonly appeared in the air (transferred as bioaerosols), but the presence of *Betsia* species (Incertae sedis; formerly Ascosphaeraceae) is suggestive of poor health of the honeybee colony. The pollen mould (*B. alvei*) is a saprophyte that lives on the pollen stored combs, especially in temperate regions [32]. Furthermore, pathogens belonging to *N. ceranae* and neogregarines were detected in the PL4 (Supplementary materials Table S2). Moreover, a high fungal spore load and a small amount of plant pollen indicates that honeybees most likely foraged on honeydew, which makes for a sugar-rich diet. At the same time, PL4 honeybees had high amounts of *γ*-proteobacteria, *Orbales, Gilliamella* (35.32%, SD=0.370) *γ-proteobacteria Rhizobiales, Bartonella* (27.06%, SD=0.560), known honeybee gut symbionts (Tab. 1A). *Gilliamella apicola* was found to be a dominant gut bacterium in honeybees and bumble bees and this bacterium simultaneously utilizes glucose, fructose, mannose, and has the ability to break down other potentially toxic carbohydrates [33].

**Table 1A.**
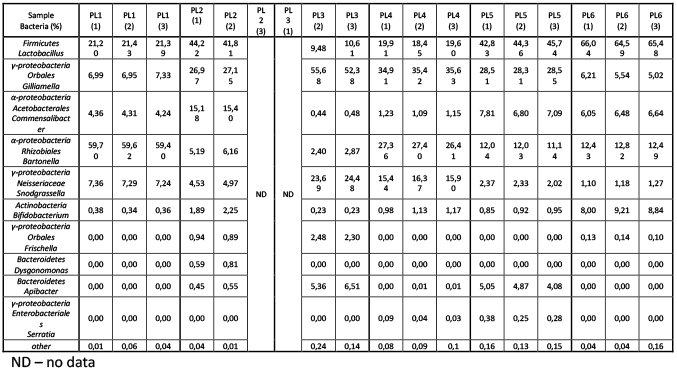
Taxonomy analysis of 16S amplicon sequencing. Six sub-groups of three bees each were analysed. Table presents relative abundance of each of the listed bacterium (numbers as percentages) in relation to the entire amplicon.

PL5 forager honeybees were collected in August and 27 taxa were identified in total from 16S amplicons analysed and 284 taxa in ITS2 (Supplementary materials Table S1). PL5 diet was mainly based on *Helianthus* sp. (34.98%, SD=1.616). PL5 had a higher number of bacteria from *Lactobacillus* (44.31%, SD=1.456) and *Gillimella* (28.46%, SD=0.129) genera (Tab. 1A). The fungal spore load was medium (59.16%, SD=2.225) and related bioaerosols (transferred as bioaerosols by wind in the air), with *Cladosporium* (11.23%, SD=2.225) and *Mycosphaerella* (6.69%, SD=1.358) as main representatives (Tab. 1B). No pathogens were detected in PL5 (Supplementary materials Table S2).

**Table 1B.**
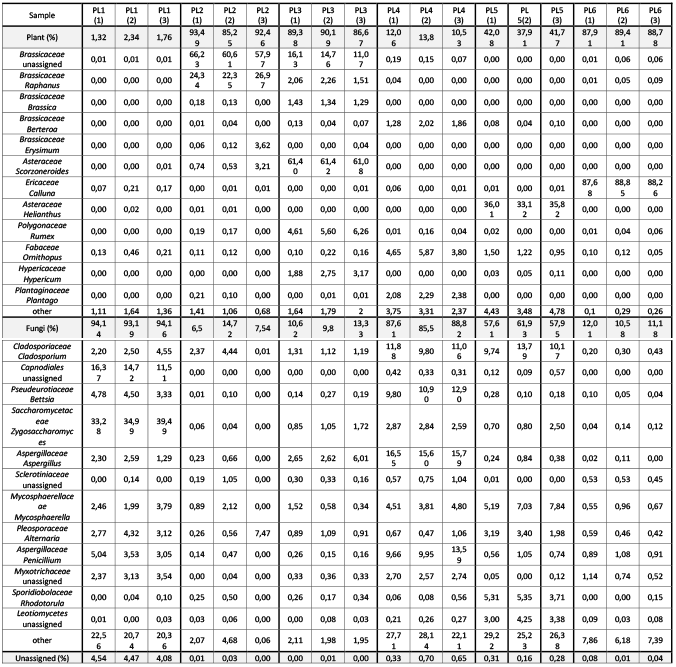
Taxonomy analysis of ITS2 amplicon sequencing. Six sub-groups of three bees each were analysed. Hits for plants and fungi were grouped into separate fractions. % for either plant or fungi is a sum of all counts for each fraction (highlighted in grey).

PL6 forager honeybees were collected in September and 28 taxa were identified in total from 16S amplicon analysed and 191 taxa in ITS2 (Supplementary materials Table S1). PL6 had the highest level of *Firmicutes, Lactobacillus* with 65.37%, SD= 0.731 and a slightly higher number of bacteria from the Comensalibacter group (Tab. 1A). PL6 honeybee diet was rich in *Calluna* pollen (88.26%, SD=0.585), which displays low protein content and is considered to be a poor source of food for bees and thus destructive for the colony development [34, 35]. The fungal spore load was moderate (11.26%, SD=0.718) with *Penicillium* as a dominant genus with 0.96%, SD=0.104 (Tab. 2). Pathogens belonging to *N. ceranae* and neogregarines were detected in PL6 (Supplementary materials Table S2).

Generally, PCA analysis of stores from spring honeybees (PL1 and PL2), summer honeybees (PL3 and PL4) and autumn honeybees (PL5 and PL6) involved splitting data into five major components which accounted for the 100% of the variation. It can be concluded from the PCA analysis that PL3 clearly differed from the others, which was mainly influenced by bacteria (b) and plants (p). PL2 and PL6 were similar to each other, bacteria (b) and plants (p) having the greatest impact as well. PL1 and PL5 were similar to each other, the greatest, albeit low influence was caused by bacteria (b) and fungi (f). PL4, similarly to PL3, clearly stood out from the others and was mainly influenced by plants (p) and fungi (f). It could also be generally seen that bacteria had the greatest influence on the variability of the plant pollen-bacteria-fungi system, followed by the plant, and then the fungi. The first two components accounted for over 53% of the variability of the entire system. Positive PC2 values may describe the summer months, and negative PC2 values to describe the months close to spring and autumn (Supplementary materials Fig. S3).

PC1 and PC3, the two main components, account for approximately 50% of the system variability (Supplementary materials Fig. S3a and S3b). It can be concluded that positive PC3 also described spring and autumn values, and negative PC3 values described the summer, which clearly differs from the others. Therefore, the third component described the season.

To summarise, we observed a prevalence of *Lactobacillus* and *Bartonella* genera in honeybees collected during spring (April PL1, May PL2) and autumn months (September PL6), while in summer months, (June PL3, July PL4, August PL5) microbiome analyses showed the prevalence of the *Gilliamella* genus, which is in agreement with previous findings^1^. In late July (PL4) the physiology of honeybees is changing due to their adaptations to overwintering and a role from two types of bacterial groups, i.e., *Lactobacilli* and *Giliamella* is suggested to be played in this process. However, increased *Lactobacilli* and *Giliamella* occurrence may simply be a consequence of the protein-rich diet. We observed fluctuations in the microbiome composition correlating with changes to the protein-richness of the pollen available in the environment. During spring and autumn, it is common for honeybees’ diet to be based on sugars ingested from honeydew. These high sugar diets can lead to fungal infections observed in April (PL1), May (PL2) and June (PL3) in honeybee samples. Probably, the detected pathogens were present in the honeybee colonies throughout the season but owing to the colony biology and well-balanced diet observed during May (PL4), June (PL3) and August (PL5), infections were less frequent among foragers and may have gone unnoticed in the whole colony screening tests.

### 2.2. Microbiome and pollen composition of honeybees from UK, Greece, Spain, and Thailand

Forager honeybees from the UK were collected on the roof of the Fogg Building at Queen Mary University of London (UK1) and in the garden of the Natural History Museum in London (UK2), in July 2019. Their bacterial microbiota was mainly composed of *Firmicutes* (*Lactobacillus*), *γ-proteobacteria* (*Orbales, Gilliamella*), *α-proteobacteria* (*Rhizobiales, Bartonella*), *γ-proteobacteria* (*Neisseriaceae, Snodgrassella*), and *Actinobacteria* (*Bifidobacterium*) (Tab. 2A). In total, 42 and 93 taxa were identified (species level) from the 16S amplicon analysis, and 70 and 94 taxa (species level) in the ITS2 in the UK1 and UK2 groups, respectively (Supplementary materials Table S1). The UK1 sample contained modest amounts of fungi (12.59%) with the prevalence of plant pollen (87.41%) derived from Apiaceae (*Ammi*), Fabaceae (*Astragalus*), Resedaceae (*Reseda*), and Fabaceae (*Styphnolobium*). UK2 honeybees foraged on Hydrangeaceae (*Hydrangea*), and Bignoniaceae plants (Tab. 2B). The UK2 sample was dominated by fungal spores (90.80%), mainly from the Myxotrichaceae, which is reported to be a common hive fungus in Europe [36, 37]. Moreover, fungal pathogen belonging to *N. ceranae* genus was detected in the UK2 (Supplementary materials Table S2).

**Table 2A.** Taxonomy analysis of 16S amplicon sequencing. Four sub-groups of two bees (UK, GR, ES) or four bees (TAI) each were analysed. Table presents relative abundance of each of the listed bacterium (numbers as percentages) in relation to the entire amplicon.

| Sample / Bacteria | UK1 | UK2 | GR1 | GR2 | ES1 | ES2 | TAI1 | TAI2 | TAI3 | TAI4 |
| --- | --- | --- | --- | --- | --- | --- | --- | --- | --- | --- |
| Country | UK | UK | Greece | Greece | Spain | Spain | Thailand | Thailand | Thailand | Thailand |
| Species | *A. mellifera* | *A. mellifera* | *A. mellifera* | *A. mellifera* | *A. mellifera* | *A. mellifera* | *A. mellifera* | *A. mellifera* | *A. cerana* | *A. cerana* |
| *Firmicutes*<br>*Lactobacillus* | 24,32 | 22,03 | 26,84 | 25,84 | 33,11 | 20,73 | 67,78 | 4,12 | 70,74 | 27,17 |
| *γ-proteobacteria*, *Orbales*<br>*Gilliamella* | 21,24 | 18,46 | 0,49 | 9,78 | 36,63 | 0,70 | 2,02 | 1,16 | 9,91 | 33,45 |
| *α-proteobacteria*, *Acetobacteriales*<br>*Commensalibacter* | 0,94 | 0,55 | 0,00 | 0,50 | 0,35 | 0,38 | 6,73 | 0,04 | 0,00 | 1,51 |
| *α-proteobacteria*, *Rhizobiales*<br>*Bartonella* | 18,51 | 0,00 | 0,14 | 49,29 | 4,53 | 66,67 | 13,30 | 0,86 | 0,00 | 9,88 |
| *γ-proteobacteria*, *Neisseriaceae*<br>*Snodgrassella* | 20,19 | 27,84 | 0,05 | 4,30 | 0,95 | 4,41 | 2,78 | 0,73 | 3,60 | 13,13 |
| *Actinobacteria*<br>*Bifidobacterium* | 7,22 | 17,54 | 54,41 | 5,55 | 8,05 | 6,21 | 4,71 | 0,27 | 0,47 | 2,63 |
| *γ-proteobacteria*, *Orbales*<br>*Frischella* | 6,13 | 6,56 | 0,00 | 0,99 | 1,58 | 0,23 | 0,00 | 0,08 | 0,00 | 8,23 |
| *Bacteroidetes*<br>*Dysgonomonas* | 0,00 | 0,00 | 0,00 | 0,00 | 5,99 | 0,00 | 0,00 | 0,00 | 0,00 | 0,00 |
| *Bacteroidetes*<br>*Apibacter* | 0,00 | 0,00 | 0,02 | 0,00 | 0,13 | 0,00 | 0,00 | 0,00 | 3,67 | 0,00 |
| *γ-proteobacteria*, *Enterobacteriales*<br>*Serratia* | 0,05 | 0,00 | 0,00 | 0,41 | 0,00 | 0,00 | 0,00 | 0,00 | 0,00 | 0,00 |
| *γ-proteobacteria*<br>*Pseudomonas* | 0,00 | 0,47 | 0,19 | 0,00 | 8,40 | 0,03 | 0,02 | 0,43 | 0,00 | 0,94 |
| *γ-proteobacteria*, *Enterobacteriales*<br>*Arsenophonus* | 0,00 | 0,00 | 0,00 | 0,00 | 0,00 | 0,00 | 0,00 | 92,11 | 0,00 | 2,42 |
| *γ-proteobacteria*, *Enterobacteriales*<br>*Klebsiella* | 0,37 | 0,00 | 0,00 | 1,43 | 0,00 | 0,00 | 0,00 | 0,00 | 0,00 | 0,00 |
| *Cyanobacteria*<br>unassigned | 0,28 | 1,78 | 11,66 | 0,01 | 0,01 | 0,00 | 0,00 | 0,13 | 0,00 | 0,15 |
| *γ-proteobacteria*,<br>*Enterobacteriaceae*<br>unassigned | 0,19 | 0,23 | 0,32 | 0,48 | 0,04 | 0,01 | 0,04 | 0,00 | 0,00 | 0,04 |
| *Actinobacteria*<br>*Cutibacterium* | 0,08 | 0,41 | 0,06 | 0,00 | 0,00 | 0,00 | 0,00 | 0,01 | 0,00 | 0,03 |
| *α-proteobacteria*, *Acetobacteriales*<br>*Bombella* | 0,03 | 0,01 | 0,00 | 0,03 | 0,04 | 0,43 | 0,10 | 0,01 | 7,05 | 0,04 |
| *Firmicutes*, *Lactobacillales*<br>*Fructobacillus* | 0,01 | 0,64 | 0,00 | 0,51 | 0,00 | 0,01 | 2,11 | 0,00 | 4,49 | 0,00 |
| *Firmicutes*, *Lactobacillales*<br>*Streptococcus* | 0,00 | 0,80 | 0,60 | 0,00 | 0,04 | 0,01 | 0,01 | 0,01 | 0,00 | 0,08 |
| *α-proteobacteria*<br>*Rickettsiales* | 0,01 | 0,05 | 4,01 | 0,00 | 0,03 | 0,00 | 0,00 | 0,04 | 0,00 | 0,12 |
| *Fusobacteria*<br>*Leptotrichia* | 0,01 | 0,25 | 0,05 | 0,00 | 0,01 | 0,00 | 0,00 | 0,00 | 0,00 | 0,00 |
| *γ-proteobacteria, Enterobacteriales*<br>*Morganella* | 0,01 | 0,00 | 0,00 | 0,43 | 0,00 | 0,00 | 0,00 | 0,00 | 0,02 | 0,00 |
| *Firmicutes*<br>*Veillonella* | 0,00 | 0,42 | 0,40 | 0,00 | 0,03 | 0,00 | 0,02 | 0,00 | 0,00 | 0,01 |
| other bacteria | 0,41 | 1,96 | 0,76 | 0,45 | 0,08 | 0,18 | 0,38 | 0,00 | 0,05 | 0,17 |

Forager honeybees from Greece (GR1, GR2) were collected in November from two colonies inhabiting the garden of the Agricultural University of Athens. In total, 80 and 31 taxa were identified from the 16S amplicon analysis, and 42 and 96 taxa in the ITS2 in the GR1 and GR2 group respectively (Supplementary materials Table S1). Although the colonies were located in a close proximity, their microbiota and food preferences varied. GR1 microbiota was mainly based on Actinobacteria (*Bifidobacterium*), Firmicutes (*Lactobacillus*), and Cyanobacteria which reached 54.41%, 26.84%, and 11.66%, respectively. Cyanobacteria indicated some colony health problems most probably connected with the contamination of water used by the honeybee colony (Tab. 2A). Moreover, pathogens belonging to *N. ceranae* and neogregarines were detected in the GR1 (Supplementary materials Table S2). The fungal load was miniscule (7.94%) and contained mainly spores present in the air transferred as bioaerosols by wind, such as Mycosphaerella and *Cladosporium*. GR1 honeybees foraged mainly on Eudicotyledonae plants. GR2 microbiome contained α-proteobacteria (Rhizobiales, *Bartonella*), Firmicutes (*Lactobacillus*), and γ-proteobacteria (Orbales, *Gilliamella*) with 49.29%, 25.84%, and 9.78%, respectively. The amount of fungal spores was miniscule (6.03%) with the dominant taxon of the Mycosphaerella reaching 1.09%. GR2 honeybees foraged mainly on Oleaceae (*Ligustrum*), Hydrangeaceae (*Hydrangea*), Myrtaceae (*Myrtus*), and Scrophulariaceae (*Buddleja*) (Tab. 2B).

**Table 2B.** Taxonomy analysis of ITS2 amplicon sequencing. Four sub-groups of two bees (UK, GR, ES) or four bees (TAI) each were analysed. Hits for plants and fungi were grouped into separate fractions. % for either plant or fungi is a sum of all counts for each fraction (highlighted in grey).

| Sample | UK1 | UK2 | GR1 | GR2 | ES1 | ES2 | TAI1 | TAI2 | TAI3 | TAI4 |
| --- | --- | --- | --- | --- | --- | --- | --- | --- | --- | --- |
| Country | UK | UK | Greece | Greece | Spain | Spain | Taiwan | Taiwan | Taiwan | Taiwan |
| Species | *A. mellifera* | *A. mellifera* | *A. mellifera* | *A. mellifera* | *A. mellifera* | *A. mellifera* | *A. mellifera* | *A. mellifera* | *A. cerana* | *A. cerana* |
| Fungi (%) | 12,58 | 90,79 | 7,96 | 6,03 | 99,87 | 99,95 | 99,28 | 7,04 | 97,75 | 99,85 |
| Saccharomycetaceae/ Lachancea | 2,44 | 0,00 | 0,00 | 0,00 | 0,00 | 0,00 | 9,09 | 0,00 | 0,18 | 0,00 |
| Aureobasidiaceae/ Aureobasidium | 0,92 | 0,45 | 0,16 | 0,07 | 0,01 | 0,00 | 0,05 | 0,00 | 0,00 | 0,35 |
| Pleosporaceae/ Alternaria | 0,67 | 1,76 | 0,63 | 0,69 | 0,04 | 0,00 | 0,35 | 0,04 | 0,00 | 0,41 |
| Cladosporiaceae / Cladosporium | 1,36 | 1,40 | 1,00 | 0,48 | 0,18 | 0,00 | 0,88 | 4,32 | 0,00 | 59,03 |
| Mycosphaerellaceae /<br>Mycosphaerella | 1,36 | 5,20 | 1,24 | 1,09 | 0,02 | 0,00 | 0,15 | 0,06 | 0,11 | 0,08 |
| Sordariaceae / unassigned | 0,00 | 0,00 | 0,00 | 0,00 | 0,00 | 0,00 | 0,00 | 0,00 | 25,37 | 0,00 |
| Botryosphaeriaceae / Lasiodiplodia | 0,00 | 0,00 | 0,00 | 0,00 | 0,00 | 0,00 | 0,00 | 0,00 | 23,78 | 0,00 |
| Aspergillaceae / Penicillium | 0,58 | 0,25 | 0,59 | 0,00 | 98,22 | 0,00 | 6,53 | 0,00 | 0,01 | 0,00 |
| Aspergillaceae / Aspergillus | 0,11 | 0,00 | 0,00 | 0,00 | 0,04 | 0,02 | 66,42 | 0,03 | 42,77 | 1,94 |
| Myxotrichaceae / unassigned | 0,03 | 73,85 | 0,00 | 0,00 | 0,00 | 0,00 | 0,00 | 0,00 | 0,00 | 0,00 |
| Saccharomycetales / Starmerella | 0,01 | 0,02 | 0,01 | 0,00 | 0,01 | 99,81 | 0,02 | 0,01 | 0,09 | 0,00 |
| Pleosporales | 0,00 | 0,13 | 0,04 | 0,04 | 0,00 | 0,00 | 0,51 | 0,79 | 0,00 | 30,38 |
| Apiosporaceae/ Arthrinium | 0,00 | 0,00 | 0,00 | 0,00 | 0,00 | 0,00 | 4,02 | 0,00 | 0,01 | 0,00 |
| other | 5,10 | 7,73 | 4,29 | 3,66 | 1,35 | 0,12 | 11,26 | 1,79 | 5,43 | 7,66 |
| Plants (%) | 87,42 | 9,21 | 92,02 | 93,97 | 0,12 | 0,04 | 0,25 | 92,94 | 0,50 | 0,15 |
| Apiaceae / Ammi | 29,78 | 0,04 | 0,00 | 0,00 | 0,00 | 0,00 | 0,00 | 0,00 | 0,00 | 0,00 |
| Fabaceae / Astragalus | 23,55 | 0,00 | 0,00 | 0,00 | 0,00 | 0,00 | 0,00 | 0,00 | 0,00 | 0,00 |
| Resedaceae / Reseda | 14,43 | 0,00 | 0,00 | 0,00 | 0,00 | 0,00 | 0,00 | 0,00 | 0,00 | 0,00 |
| Fabaceae / Styphnolobium | 12,94 | 0,00 | 0,00 | 0,00 | 0,00 | 0,00 | 0,00 | 0,00 | 0,00 | 0,00 |
| Hypericaceae / Hypericum | 2,95 | 0,00 | 0,00 | 0,62 | 0,00 | 0,00 | 0,00 | 0,00 | 0,00 | 0,00 |
| Oleaceae / Ligustrum | 0,43 | 0,20 | 0,08 | 22,39 | 0,00 | 0,00 | 0,00 | 0,00 | 0,00 | 0,00 |
| Hydrangeaceae / Hydrangea | 0,31 | 1,22 | 0,51 | 38,51 | 0,00 | 0,00 | 0,00 | 0,00 | 0,00 | 0,00 |
| Bignoniaceae / unassigned | 0,00 | 5,58 | 0,00 | 0,00 | 0,00 | 0,00 | 0,00 | 0,00 | 0,00 | 0,00 |
| Myrtaceae / Myrtus | 0,00 | 0,85 | 0,00 | 7,96 | 0,00 | 0,00 | 0,00 | 0,00 | 0,00 | 0,00 |
| Scrophulariaceae / Buddleja | 0,00 | 0,12 | 0,12 | 8,07 | 0,00 | 0,00 | 0,00 | 0,00 | 0,00 | 0,00 |
| Asteraceae / unassigned | 0,00 | 0,09 | 0,00 | 0,00 | 0,00 | 0,00 | 0,00 | 91,33 | 0,00 | 0,00 |
| Asteraceae / Hypochaeris | 0,00 | 0,04 | 0,00 | 2,35 | 0,00 | 0,00 | 0,00 | 0,00 | 0,00 | 0,00 |
| Eudicotyledonae / unassigned | 0,00 | 0,00 | 89,14 | 0,00 | 0,00 | 0,00 | 0,00 | 0,00 | 0,00 | 0,00 |
| Myrtaceae / Eucalyptus | 0,00 | 0,00 | 1,49 | 0,44 | 0,00 | 0,00 | 0,00 | 0,00 | 0,00 | 0,00 |
| Cucurbitaceae / Lagenaria | 0,00 | 0,00 | 0,00 | 3,45 | 0,00 | 0,00 | 0,00 | 0,00 | 0,00 | 0,00 |
| Asteraceae / Picris | 0,00 | 0,00 | 0,00 | 2,31 | 0,00 | 0,00 | 0,00 | 0,00 | 0,00 | 0,00 |
| Fabaceae / Pterocarpus | 0,00 | 0,00 | 0,00 | 0,00 | 0,00 | 0,00 | 0,16 | 0,00 | 0,42 | 0,00 |
| Fabaceae / Mimosa | 0,00 | 0,00 | 0,00 | 0,00 | 0,00 | 0,00 | 0,08 | 0,67 | 0,00 | 0,15 |
| Asteraceae / Bidens | 0,00 | 0,00 | 0,00 | 0,00 | 0,00 | 0,00 | 0,00 | 0,65 | 0,00 | 0,00 |
| Myrtaceae / Syzygium | 0,00 | 0,00 | 0,00 | 0,00 | 0,00 | 0,00 | 0,00 | 0,00 | 0,06 | 0,00 |
| Araliaceae / Hedera | 0,00 | 0,00 | 0,07 | 0,00 | 0,12 | 0,04 | 0,00 | 0,00 | 0,00 | 0,00 |
| other | 3,03 | 1,07 | 0,61 | 7,87 | 0,00 | 0,00 | 0,01 | 0,29 | 0,02 | 0,00 |
| Unassigned | 0,00 | 0,00 | 0,02 | 0,00 | 0,01 | 0,01 | 0,47 | 0,02 | 1,75 | 0,00 |

Forager honeybees from Spain (ES1, ES2) were collected in November from experimental colonies located near Marchamalo. In total, 34 and 25 taxa were identified from the 16S amplicon analysis, and 31 and 13 taxa in the ITS2 in the ES1 and ES2 groups, respectively (Supplementary materials Table S1). ES1 contained γ-proteobacteria (Orbales, *Gilliamella*), and Firmicutes (*Lactobacillus*), at the level of 36.63%, and 33.11%, respectively (Tab. 2A). These honeybees most likely foraged on honeydew since pollen DNA was hardly detected (Araliaceae *(Hedera)* 0.12% for ES1 and 0.04% for ES2). ES1 and ES2 had dominant fungal fraction containing spores transferred as bioaerosols by wind in the air, such as *Penicillium*, *Cladosporium* and *Mycosphaerella* (Tab. 2B). Additionally, pathogens belonging to *N. ceranae* and neogregarines were detected in the ES2 (Supplementary materials Table S2).

Forager honeybees from Thailand consisted of both: western honeybee (*Apis mellifera* TAI1 and TAI2) and Asian honeybee (*Apis cerana* TAI3 and TAI4). In eastern Asia, these two bee genera inhabit the same locations resulting in the transfer of pathogens from *Apis cerana* to *Apis mellifera*, as described for *Varroa destructor* and *Nosema ceranae*. Thai samples were collected in February, the best month in the year for honeybee colonies in Thailand. In total, 29, 22, 19 and 42 taxa were identified from the 16S amplicon analysis, and 50, 94, 59 and 85 taxa in the ITS2 in the TAI1, TAI2, TAI3 and TAI4 groups, respectively (Supplementary materials Table S1). TAI1 contained high loads of Firmicutes (*Lactobacillus*) and *Bartonella* (Tab. 2A), and a high quantity of fungal spores such as *Aspergillus* (66.42%), Saccharomycetaceae (9.09%), and *Peniclilium* (6.53%) (Tab. 2B). A trace amount of plant pollen was detected in TAI1 *(Pterocarpus* with 0.16%, *Mimosa* 0.08%) indicating honeydew to have been the main source of forage (Tab. 2B). Bacterial microbiota of TAI2 was mainly composed of *Arsenophonus* bacteria (γ-proteobacteria, Enterobacteriales) 92.11% of which were insects’ intracellular symbionts. *Arsenophonus* species showed a broad spectrum of symbiotic relationships varying from parasitic son-killers to coevolving mutualists [37]. Moreover, pathogens belonging to *N. ceranae* and neogregarines were detected in TAI1 (Supplementary materials Table S2). TAI2 honeybees foraged mainly on Asteraceae pollen (91.33%) and contained only a miniscule amount of fungal spores transferred as bioaerosols by wind in the air, as *Cladosporium* 4.32%. *Apis cerana* samples contained 70.74% of *Lactobacillus* genus for TAI3, 27.17% for TAI4, *Gilliamella*, 33.45% for TAI4, and *Snodgrassella genus* with 3.60% and 13.13% for TAI3 and TAI4, respectively. Fungal spores present in the samples were related with the air bioaerosols, including *Aspergillus* 42.77% for TAI3, *Cladosporium* 59.03% and *Pleosporales* 30.38% for TAI4. The amount of plant pollen was miniscule (*Pterocarpus* with 0.42% for TAI3, *Mimosa* 0.15% for TAI4) indicating honeydew as the main forage for Thai *A. cerana* bees (Tab. 2). Moreover, pathogens belonging to *N. ceranae* and neogregarines were detected in TAI4 (Supplementary materials Table S2). PCA analysis of the honeybees’ stores split data into five major components which accounted for the 100% of the variation. PC1 and PC2 components accounted for nearly 57% of the variation, respectively 32.83% and 23.65 (Fig. 3, figures a and b should be considered simultaneously).

The PCA analysis allowed us to determine the differences between bees from different countries. Four different areas were distinguished. Clear differences can be seen between bee samples from Greece, UK, Spain, Poland, and the two samples of bees from Thailand (Fig. 3 Figure b).

**Figure 3.**
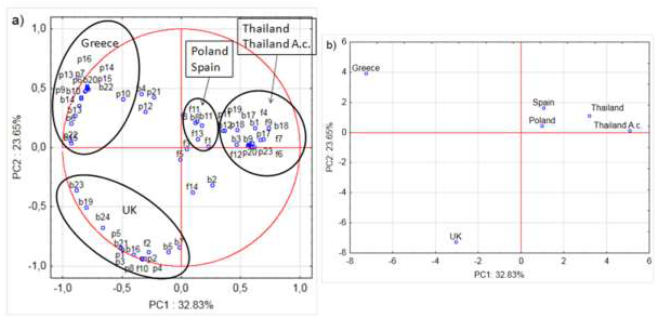
Loading plot (a) and score plot (b) of the principal components’ analysis (PC1 and PC2) carried out on the analytical data of the taxonomy detected in the world bees (Thailand A.c. - *A. cerana*). Small letters on loading plot (a): **b** – data obtained from **bacteria** NGS analysis, **f** – data obtained from **fungal** NGS analysis, **p** - data obtained from **plant** NGS analysis.

PCA analysis shows Greek samples to markedly differ from the others, which is mainly influenced by bacteria (b) and plants (p). The PCA analysis findings for UK bee samples also clearly differ from the others which is mainly due to bacteria (b), then plants (p) and finally fungi (f). One can also distinguish samples from Spain and Poland from the other samples, which is mostly influenced by bacteria (b) and fungi (f), however, these are similar to the results for the samples from Thailand which are mostly influenced by taxons of bacteria (b), plants (p) and fungi (f). Taxons originating from bacteria, and also, to a lesser extent, those originating from plants and fungi can be said to have the greatest influence on the variability of the system (Fig. 3a and 3b).

## 3. Conclusions

Foragers are worker honeybees with an age of over 21 days, at which time they collect water, nectar and pollen as well as supplements necessary for the colony to survive. All forager bees are in an comparable age, have similar function and physiological processes [39] and similar microbiota. Therefore, they should share similar microbiota. However, studies indicated that forager honeybees have “contingent microbiome” dependent mainly of food they forage [2, 40, 41]. This carries a danger, because with poor food resources, the microbiota will be inappropriate and non-functioning [42]. Nowadays many factors influence bee microbiota e.g.: monocultures, nutritional stress, pesticide exposure, agrochemicals, many of which exhibit hidden antimicrobial properties contributing greatly to honeybee stress tolerance and disease resistance, which leads to higher honeybee mortality and a high rate of colony loss [5, 43], pathogens which trigger bee malnutrition [44], changes in the composition of their microelements [45] and yeast content [46]. Honeybee dietary preferences, originally developed during the course of evolution, might not presently be favourable for honeybees. It is an urgent issue that should be carefully studied to aid bees to survive in the anthropogenic biosphere.

Pollination is a crucial process for the maintenance of plant-based food supplies [47]. To maintain the health of honeybees, it is favourable to prevent the spread of a disease, prevent exposure to insecticides and pesticides and provide a variety of plant pool to maintain optimal nutrition and microbiome throughout the season [48]. The use of NGS techniques in the identification of the pollen pool preferentially chosen by honeybees can provide strategies to maintain healthy colonies of bees. This technique can greatly expand and supplement the knowledge based on long term observation for the most efficient “pollinator-friendly” plants [49, 50]. This should help to create more effective pollinator beneficial plant species composition for “pollinator-friendly” gardens or to enrich the plant species on flower strips.

In our research, the composition of honeybee microbiomes was mainly determined by dietary preferences. Even when colonies originating from one apiary, for instance from Spain or Greece, honeybees chose different plants to forage on. UK bees from a single highly urbanised area (London) exhibited particularly high diversity, chose different food sources and were otherwise prone to diseases. Furthermore, honeybees choosing honeydew-rich died were more susceptible to pathogens (*Nosema ceranae* and neogregarines). The period when honeybees switch to the winter generation (longer-lived forager honeybees) was the time when the whole colony proved most sensitive to dietary perturbations.

Our findings are in line with limited other reports, that suggested honeybees from varying apiaries take independent decisions on the choice of pollen and nectar they forage upon [51]. As observed, colonies of bees located in close proximities may have varying pollen composition in their gut. It remains unclear how bees decide what pollen to forage. However, flower structure, nectar volume, sugar content and composition were indicated to play a role in attracting bees [34, 35, 49]. Some studies indicated that honeybees primarily chose pollen rich in essential amino acids [34, 35, 49]. On the other hand, honeybees affected by diseases because of their considerable energy need and having being weakened by diseases, were unable to undertake forage flights to collect good quality pollen and collected one which had worse nutritional parameters. It is possible that evolutionary adaptation of bees fails to benefit them in the modern anthropomorphised environment.

## 4. Highlights

1. Honeybees were more susceptible to pathogens if they did not receive a well-balanced diet and honeybees with a honeydew diet were more prone to fungal pathogens (*Nosema ceranae*) and neogregarines.
2. The period when honeybees switch to the winter generation (longer-lived forager honeybees) is the most sensitive to diet perturbations and hence pathogens attack, for the whole beekeeping season.
3. The composition of honeybee microbiomes is mainly determined by dietary preferences.

## 4. Materials and Methods

### 5.1. Honeybee collection and DNA isolation

Forager honeybees were collected from one location in Lublin, Poland [[51°15′N_22°34′E](https://geohack.toolforge.org/geohack.php?pagename=Lublin&params=51_14_53_N_22_34_13_E_region:PL_type:city)] each month from April to September 2018 (PL1-PL6). Forager honeybees from UK were collected from the roof of the Fogg Building of Queen Mary University of London (UK2) [[51°52’N_0°03’W](https://geohack.toolforge.org/geohack.php?pagename=Mile_End&params=51.5248_N_0.0314_W_region:GB_type:city)] and in the garden of the Natural History Museum in London (UK1) [[51°29′N_0°10′W](https://tools.wmflabs.org/geohack/geohack.php?language=pl&pagename=Muzeum_Historii_Naturalnej_w_Londynie&params=51.495833333333_N_0.17611111111111_W)] in July 2019. Greek (GR1, GR2) samples of forager honeybees were collected in November 2017 from two colonies inhabiting the garden of The Agricultural University of Athens [[37°59′N_23°42′E](https://tools.wmflabs.org/geohack/geohack.php?language=pl&pagename=Atenski_Uniwersytet_Rolniczy&params=37.983888888889_N_23.705555555556_E)]. Forager honeybees from Spain (ES1, ES2) were collected in November 2017 from experimental colonies located at Marchamalo (Centro Apícola y Agroambiental de Marchamalo (CIAPA-IRIAF), Marchamalo, Spain [[40°68′N_3°21′W](https://tools.wmflabs.org/geohack/geohack.php?language=pl&pagename=Marchamalo&params=40.668611111111_N_3.2019444444444_W)]. Thai samples of forager bees consisting of both western honeybee (*Apis mellifera*) and Asian honeybee (*Apis ceranae*) were collected in February 2018 in the proximity of Chiang Mai University [[18°50′_98°58″E](https://geohack.toolforge.org/geohack.php?pagename=Chiang_Mai_Province&params=18_50_14_N_98_58_14_E_region:TH_type:adm1st_dim:200000_source:dewiki)], *Apis mellifera* samples were marked TAI1 and TAI2, and *A. ceranae* as TAI3 and TAI4. Genomic DNA was extracted from whole honeybees using QIAamp DNA Kit according to manufacturer’s instructions. Isolates were sent to the Biobank, Poland for NGS analysis.

### 5.2. NGS

NGS sequencing and the analysis of the 16S rRNA bacterial gene amplicon was based on the V3-V4 region and the ITS2 eukaryotic region for bee DNA samples. Amplicon libraries, were prepared using the *16S Metagenomic Sequencing Library Preparation, Preparing 16S Ribosomal RNA Gene Amplicons for the Illumina MiSeq System* (Illumina^®^) protocol. Information about primers sequences, PCR conditions is shown in Supplementary materials Table S3.

All data are available at https://www.ncbi.nlm.nih.gov/bioproject/PRJNA686953.

### 5.3. Positive, negative control

The positive qualitative control for the V3-V4 region of the 16s rRNA gene was the DNA isolate derived from an ear swab. For the ITS2 region, it was DNA isolated from the *Saccharomyces cerevisiae* strain. PCR grade water was the negative qualitative control for both kinds of amplicons.

### 5.4. Purification, clean-up

The amplicons obtained were purified using magnetic beads (AMPure XP beads; Beckman Coulter) according to Illumina^^®^^ protocol.

### 5.5. Library pooling – concentration, normalization

Before pooling samples for libraries, the concentration was measured. The concentration [ng/uL] was measured using the NanoDrop™ 2000/c Spectrophotometer (Thermo Fisher Scientific) for each amplicon. Samples were diluted (PCR grade water) to the same concentration and pooled. To determine the final library concentration in [nM], the *NEBNext Ultra DNA Library Prep Kit for Illumina* (New England Biolabs^®^ Inc.) protocol was followed. The final concentration of pooled libraries for sequencing was 8 pM.

### 5.6. Sequencing

Prepared libraries were sequenced on an Illumina MiSeq platform, 2 x 300 sequence reading in paired ends mode. The run contained PhiX libraries (PhiX Control Kit v3, Illumina^®^), to serve as an internal positive quality control.

### 5.7. 16S rRNA bacterial gene analyses

Reads from the sequencing run were imported into the QIIME 2 version 2019.10 artifact [52]. Then sequences were trimmed at first 21 bp for forward and reverse reads and truncated to 250 for forward reads and 240 for reverse reads. Reads were then denoised with DADA2 [53] and merged together. Sequences were aligned with MAFFT [54] and used to construct a phylogeny with fasttree [55]. Rarefaction was performed with at least 14096 sequences per sample for subsequent stages of the analysis. Taxonomic assignments of representative sequences were conducted using q2-feature-classifìer with the sklearn classifier [56] trained on SILVA 132 database at 99% similarity level [57].

### 5.8. ITS2 region analyses

Analogous steps as for “*16S rRNA bacterial gene analyses*” were performed for the ITS2 analysis. Reads were trimmed and denoised separately, then they were merged for further analysis. Then data were trimmed at first 21 bp for forward and reverse reads and truncated to 300 for forward reads and 215 for reverse reads. Taxonomic assignments were conducted analogous to 16S analysis with classifier trained on ITS gene clustered at 99% similarities within UNITE database released 04.02.2020 containing all eukaryotes [58]. Sampling depth was set to 34100 sequences for the diversity analyses.

### 5.9. Analyses of amplicon from honeybee intestine samples

Amplicons for the 16S region and ITS2 (Supplementary materials Table S4) were sequenced using the Illumina MiSeq platform. Data were trimmed and merged. For 16S analyses only full-length reads over 229 bp with medium length of all sequences at 414 bp were used. Sequences were assigned to taxonomy using classifier trained on SILVA 132 database with minimum similarity 90% of read matching to the reference. For ITS2 analyses only full-length reads over 269 bp with medium length of all sequences at 337 bp were used. Sequences were assigned to taxonomy using classifier trained on all eukaryotes UNITE database v8.2 with the minimum similarity of 90% of the read matching to the reference (Supplementary materials Table S4 in Excel).

### 5.10. Screening for pathogen infected honeybee samples

Isolated DNA was used as the template for screening pathogens: *Nosema apis*, *Nosema ceranae*, *Nosema bombi*, tracheal mite (*Acarapis woodi*), any organism in the parasitic order Trypanosomatida, including Crithidia spp. (i.e., *Crithidia mellificae)*, neogregarines including *Mattesia* and *Apicystis* spp. (i.e., *Apicistis bombi)*, using PCR techniques described earlier [59–62]. Primers used for pathogen detection are listed in the Supplementary materials Table S2. Detection of the pathogens in honeybee samples.

### 5.11. Statistical analysis

Analyses of correlations and Principal Component Analysis (PCA) were performed using software Statistica (version 12.0, StatSoft Inc., USA) at the significance level of α = 0.05. The analysis was used to determine the relationships between the bee sample and the bacterial group, plant group, and fungi group. The optimum number of principal components obtained in the PCA analysis was established based on Cattell’s criterion. The data matrix for the PCA of the Polish bees had 37 columns and 6 rows and of the world samples of bees had 61 columns and 6 rows (UK, Spain, Greek, Thailand and Poland) had 61 columns and 6 rows. The input matrix was auto scaled.

## Acknowledgments

The publication of the article was financed by the Polish National Agency for Academic Exchange under the Foreign Promotion Programme (NAWA), for AAP (bee-research.umcs.pl; Api Lab UMCS PPI/PZA/2019/1/00039). Honeybees were collected during the Miniatura 2 project ID 418332 founded by the NCN for AAP and the first research plans were possible thanks to EU: GB-TAF-7137 SYNTHESYS project for AAP. Work in the Starosta lab was funded by EMBO Installation Grants 2017, and POIR. 04.04.00-00-3E9C/17-00 for ALS. Work in the Hurd lab was funded by the BBSRC (BB/L023164/1) and granted to PJH.

## Author Contributions

AAP (senior author), designed the experiments, analysed data, and wrote the paper. PL, analysed data, prepared figures, supplemental information, and methods. PH, analysed data, especially of metabiom and parasites, co-wrote the paper. AP, analysed UK data. JMM, conducted laboratory work for sequencing library preparation, sequencing, and detection of pathogens. ŁG, analysed raw data from metabiom sequencing, prepared tables, co-wrote the paper. DS, analysed data from metabiom sequencing, co-wrote the paper. SG, performed genetic analyses. DZ, drafted and made a correction of the manuscript. MG, and RR, analysed data using PCA, interpreted the results, co-wrote the paper. PK, analysed Thai data. RMH, and MH, analysed UK data, co-wrote the paper. ALS, analysed data, prepared figures and tables and co-wrote the paper. All authors read and approved the final manuscript.

## Lead Contact

Further information and requests for resources and reagents should be directed to and will be fulfilled by the Lead Contact, Aneta A. Ptaszyńska.

---

**Table 1.** Taxonomy analysis of 16S and ITS amplicon sequencing. A full list of all detected organisms is listed in Supplemental Table S4.

## References

1. Yao, R., Xu, L., Lu, G., & Zhu, L. (2019). Evaluation of the function of wild animal gut microbiomes using next-generation sequencing and bioinformatics and its relevance to animal conservation. Evolutionary Bioinformatics, 15, 1176934319848438. https://doi.org/10.1177/1176934319848438

2. Romero, S., Nastasa, A., Chapman, A., Kwong, W.K., and Foster, L.J. (2019). The honey bee gut microbiota: strategies for study and characterization. Insect Mol. Biol. 28, 455–472. Available at: http://doi.wiley.com/10.1111/imb.12567.

3. Hawkins J, de Vere N, Griffith A, Ford CR, Allainguillaume J, Hegarty MJ, et al. (2015) Using DNA Metabarcoding to Identify the Floral Composition of Honey: A New Tool for Investigating Honey Bee Foraging Preferences. PLoS ONE 10(8): e0134735. https://doi.org/10.1371/journal.pone.0134735

4. Kešnerová, L., Emery, O., Troilo, M., Liberti, J., Erkosar, B., and Engel, P. (2020). Gut microbiota structure differs between honeybees in winter and summer. ISME J. 14, 801–814. Available at: https://pubmed.ncbi.nlm.nih.gov/31836840/.

5. Tauber, J.P., Nguyen, V., Lopez, D., and Evans, J.D. (2019). Effects of a resident yeast from the honeybee gut on immunity, microbiota, and Nosema disease. Insects 10. Available at: https://pubmed.ncbi.nlm.nih.gov/31540209/.

6. McNeil, D.J., McCormick, E., Heimann, A.C. et al. Bumble bees in landscapes with abundant floral resources have lower pathogen loads. Sci Rep 10, 22306 (2020). https://doi.org/10.1038/s41598-020-78119-2

7. Patel, V., Pauli, N., Biggs, E. et al. Why bees are critical for achieving sustainable development. Ambio 50, 49–59 (2021). https://doi.org/10.1007/s13280-020-01333-9

8. Zalasiewicz, J., Waters, C. and Williams, M. Chapter 32: The Anthropocene. In: Gradstein, F., Ogg, J., Schmitz, M. and Ogg, G. (eds.) A Geologic Time Scale 2020.

9. Howard J. (2017) Anthropogenic Landforms and Soil Parent Materials. In: Anthropogenic Soils. Progress in Soil Science. Springer, Cham. https://doi.org/10.1007/978-3-319-54331-4_3

10. Le Conte Y, Navajas M. Climate change: impact on honey bee populations and diseases. Rev Sci Tech. 2008 Aug;27(2):485–97, 499-510. English, French. PMID: 18819674.

11. IUCN Red List, IUCN Species Survival Commission. Guidelines for Application of IUCN Red List criteria at regional and national levels. Table 3; Last Updated: Dec. 2020 List version 2020–3 https://www.iucnredlist.org/statistics

12. Hallmann, C. A., Sorg, M., Jongejans, E., Siepel, H., Hofland, N., Schwan, H. et al., More than 75 percent decline over 27 years in total flying insect biomass in protected areas. PLoS One 12, e0185809 (2017).

13. Fox, R. The decline of moths in Great Britain: a review of possible causes. Insect Conserv. Divers. 6, 5–19 (2013).

14. Benton, T. G., Bryant, D. M., Cole, L. & Crick, H. Q. Linking agricultural practice to insect and bird populations: a historical study over three decades. J. Appl. Ecol. 39, 673–687 (2002).

15. Thomas, J. A., Telfer, M. G., Roy, D. B., Preston, C. D., Greenwood, J., Asher, J. et al., Comparative losses of British butterflies, birds, and plants and the global extinction crisis. Science 303, 1879–1881 (2004).

16. Hallmann, C. A., Foppen, R. P., van Turnhout, C. A., de Kroon, H. & Jongejans, E. Declines in insectivorous birds are associated with high neonicotinoid concentrations. Nature 511, 341–343 (2014).

17. Pimm, S. L., Jenkins, C. N., Abell, R., Brooks, T. M., Gittleman, J. L., Joppa, L. N., Raven, P. H., Roberts, C. M. & Sexton, J. O. The biodiversity of species and their rates of extinction, distribution, and protection. Science 344, 1246752 (2014).

18. Sánchez-Bayo, F. & Wyckhuys, K. A. G. Worldwide decline of the entomofauna: A review of its drivers. Biol. Conservation 232, 8–27 (2019).

19. Marshman, J.; Blay-Palmer, A.; Landman, K. Anthropocene Crisis: Climate Change, Pollinators, and Food Security. Environments 2019, 6, 22.

20. EU Pollinators Initiative. Brussels, 1.6.2018. COM(2018) 395 final. https://eur-lex.europa.eu/legal-content/EN/TXT/PDF/?uri=CELEX:52018DC0395&from=PL

21. “LIFE+ URBANBE” https://ec.europa.eu/environment/life/project/Projects/index.cfm?fuseaction=home.showFile&rep=file&fil=URBANBEES_Management_Plan.pdf

22. “City Bees” https://www.eea.europa.eu/atlas/eea/city-bees/story/article,

23. “Urban Beekeeping” https://beeproject.ca/urban-beekeeping

24. Ellegaard, K.M., and Engel, P. (2019). Genomic diversity landscape of the honey bee gut microbiota. Nat. Commun. 10, 1–13. Available at: https://doi.org/10.1038/s41467-019-08303-0.

25. Ahn, J.H., Hong, I.P., Bok, J.I., Kim, B.Y., Song, J., and Weon, H.Y. (2012). Pyrosequencing analysis of the bacterial communities in the guts of honey bees Apis cerana and Apis mellifera in Korea. J. Microbiol. 50, 735–745. Available at: http://www.springerlink.com/content/120956.

26. Cilia G., Fratini F., Tafi E., Mancini S., Turchi B., Sagona S., Cerri D., Felicioli A. and Nanetti A. (2021) Changes of Western honey bee Apis mellifera ligustica (Spinola, 1806) ventriculus microbial profile related to their in-hive tasks, Journal of Apicultural Research, 60:1, 198–202, DOI: 10.1080/00218839.2020.1830259

27. Laha, R.C., De Mandal, S., Ralte, L., Ralte, L., Kumar, N.S., Gurusubramanian, G., Satishkumar, R., Mugasimangalam, R., and Kuravadi, N.A. (2017). Meta-barcoding in combination with palynological inference is a potent diagnostic marker for honey floral composition. AMB Express 7, 132.

28. Gous, A., Swanevelder, D.Z.H., Eardley, C.D., and Willows-Munro, S. (2019). Plant-pollinator interactions over time: Pollen metabarcoding from bees in a historic collection. Evol. Appl. 12, 187–197. Available at: http://doi.wiley.com/10.1111/eva.12707.

29. Keller, A., Danner, N., Grimmer, G., Ankenbrand, M., von der Ohe, K., von der Ohe, W., Rost, S., Härtel, S., and Steffan-Dewenter, I. (2015). Evaluating multiplexed next-generation sequencing as a method in palynology for mixed pollen samples. Plant Biol. 17, 558–566. Available at: https://pubmed.ncbi.nlm.nih.gov/25270225/.

30. Baksay, S., Pornon, A., Burrus, M., Mariette, J., Andalo, C., and Escaravage, N. (2020). Experimental quantification of pollen with DNA metabarcoding using ITS1 and trnL. Sci. Rep. 10, 1–9. Available at: https://doi.org/10.1038/s41598-020-61198-6.

31. Szczęsna T. 2006. Protein content and amino acid composition of bee-collected pollen from selected botanical origins. Journal of Apicultural Science. Volume 50, Issue 2, Pages 81–90.

32. Wynns, A.A. Convergent evolution of highly reduced fruiting bodies in Pezizomycotina suggests key adaptations to the bee habitat. BMC Evol Biol 15, 145 (2015). https://doi.org/10.1186/s12862-015-0401-6

33. Zheng, H., Nishida, A., Kwong, W. K., Koch, H., Engel, P., Steele, M. I., & Moran, N. A. (2016). Metabolism of Toxic Sugars by Strains of the Bee Gut Symbiont Gilliamella apicola. mBio, 7(6), e01326–16. https://doi.org/10.1128/mBio.01326-16

34. Vanderplanck, M., Vereecken, N., Grumiau, L. et al. The importance of pollen chemistry in evolutionary host shifts of bees. Sci Rep 7, 43058 (2017). https://doi.org/10.1038/srep43058

35. Vanderplanck M, Moerman R, Rasmont P, Lognay G, Wathelet B, et al. (2014) How Does Pollen Chemistry Impact Development and Feeding Behaviourof Polylectic Bees? PLoS ONE 9(1): e86209. doi:10.1371/journal.pone.0086209

36. Rodríguez-Andrade, E., Stchigel, A.M., Terrab, A. et al. Diversity of xerotolerant and xerophilic fungi in honey. IMA Fungus 10, 20 (2019). https://doi.org/10.1186/s43008-019-0021-7

37. Burnside CE (1929) Saprophytic fungi associated with the honey bee. Bee World 10:42. https://doi.org/10.1080/0005772X.1929.11092782

38. Nováková, E., Hypša, V. & Moran, N.A. *Arsenophonus*, an emerging clade of intracellular symbionts with a broad host distribution. BMC Microbiol 9, 143 (2009). https://doi.org/10.1186/1471-2180-9-143

39. Abou-Shaara, H.F. (2014). The foraging behaviour of honey bees, Apis mellifera: A review. Vet. Med. (Praha). 59, 1–10.

40. Khan, K.A., Al-Ghamdi, A.A., Ghramh, H.A., Ansari, M.J., Ali, H., Alamri, S.A., Al-Kahtani, S.N., Adgaba, N., Qasim, M., and Hafeez, M. (2020). Structural diversity and functional variability of gut microbial communities associated with honey bees. Microb. Pathog. 138, 103793.

41. Kwong, W.K., and Moran, N.A. (2016). Gut microbial communities of social bees. Nat. Rev. Microbiol. 14, 374–384. Available at: https://www.nature.com/articles/nrmicro.2016.43.

42. Daisley, B.A., Chmiel, J.A., Pitek, A.P., Thompson, G.J., and Reid, G. (2020). Missing Microbes in Bees: How Systematic Depletion of Key Symbionts Erodes Immunity. Trends Microbiol.

43. Kakumanu, M.L., Reeves, A.M., Anderson, T.D., Rodrigues, R.R., and Williams, M.A. (2016). Honey Bee Gut Microbiome Is Altered by In-Hive Pesticide Exposures. Front. Microbiol. 7, 1255.

44. Ptaszyńska A.A., Borsuk G., Mułenko W., Demetraki-Paleolog J. (2014). Differentiation of Nosema apis and Nosema ceranae spores under Scanning Electron Microscopy (SEM). Journal of Apicultural Research, 53, 537–544, DOI: 10.3896/IBRA.1.53.5.02

45. Ptaszyńska, A.A.; Gancarz, M.; Hurd, P.J.; Borsuk, G.; Wiaćcek, D.; Nawrocka, A.; Strachecka, A.; Załuski, D.; Paleolog, J. Changes in the bioelement content of summer and winter western honeybees (Apis mellifera) induced by Nosema ceranae infection (2018). PLoS ONE, 13, e0200410.

46. Ptaszyńska A.A., Paleolog J., Borsuk G. 2016. Nosema ceranae Infection Promotes Proliferation of Yeasts in Honey Bee Intestines. PLoS ONE 11(10): e0164477. doi:10.1371/journal.pone.0164477

47. Costanza, R., D’Arge, R., De Groot, R., Farber, S., Grasso, M., Hannon, B., Limburg, K., Naeem, S., O’Neill, R. V., Paruelo, J., et al. (1997). The value of the world’s ecosystem services and natural capital. Nature 387, 253–260.

48. Goulson, D., Nicholls, E., Botias, C., and Rotheray, E.L. (2015). Bee declines driven by combined stress from parasites, pesticides, and lack of flowers. Science (80-.). 347, 1255957–1255957.

49. Mayer, C., Adler, L., Armbruster, S., Dafni, A., Eardley, C., Huang, S.-Q., Kevan, P.G., Ollerton, J., Packer, L., Ssymank, A., et al. (2011). Pollination ecology in the 21st Century: Key questions for future research. J. Pollinat. Ecol. 3, 8–23.

50. De Vere, N., Jones, L.E., Gilmore, T., Moscrop, J., Lowe, A., Smith, D., Hegarty, M.J., Creer, S., and Ford, C.R. (2017). Using DNA metabarcoding to investigate honey bee foraging reveals limited flower use despite high floral availability. Sci. Rep. 7, 1–10. Available at: https://www.nature.com/scientificreports.

51. Vaudo, A.D., Tooker, J.F., Grozinger, C.M., and Patch, H.M. (2015). Bee nutrition and floral resource restoration. Curr. Opin. Insect Sci. 10, 133–141.

52. Bolyen, E, Rideout, J, Dillon, M, Bokulich, N, Abnet, C, Al-Ghalith, G, Alexander, H, Alm, E, Arumugam, M, Asnicar, F, Bai, Y, Bisanz, J, Bittinger, K, Brejnrod, A, Brislawn, C, Brown, C, Callahan, B, Caraballo-Rodríguez, A, Chase, J, Cope, E, Da Silva, R, Diener, C, Dorrestein, P, Douglas, G, Durall, D, Duvallet, C, Edwardson, C, Ernst, M, Estaki, M, Fouquier, J, Gauglitz, J, Gibbons, S, Gibson, D, Gonzalez, A, Gorlick, K, Guo, J, Hillmann, B, Holmes, S, Holste, H, Huttenhower, C, Huttley, G, Janssen, S, Jarmusch, A, Jiang, L, Kaehler, B, Kang, K, Keefe, C, Keim, P, Kelley, S, Knights, D, Koester, I, Kosciolek, T, Kreps, J, Langille, M, Lee, J, Ley, R, Liu, YX, Loftfield, E, Lozupone, C, Maher, M, Marotz, C, Martin, B, McDonald, D, McIver, L, Melnik, A, Metcalf, J, Morgan, S, Morton, J, Naimey, A, Navas-Molina, J, Nothias, L, Orchanian, S, Pearson, T, Peoples, S, Petras, D, Preuss, M, Pruesse, E, Rasmussen, L, Rivers, A, Robeson, M, Rosenthal, P, Segata, N, Shaffer, M, Shiffer, A, Sinha, R, Song, S, Spear, J, Swafford, A, Thompson, L, Torres, P, Trinh, P, Tripathi, A, Turnbaugh, P, Ul-Hasan, S, Hooft, J, Vargas, F, Vázquez-Baeza, Y, Vogtmann, E, Hippel, M, Walters, W, Wan, Y, Wang, M, Warren, J, Weber, K, Williamson, C, Willis, A, Xu, Z, Zaneveld, J, Zhang, Y, Zhu, Q, Knight, R, Caporaso, J. “Reproducible, interactive, scalable and extensible microbiome data science using QIIME 2”. Nature Biotechnology 2019; 37(8):852–857.

53. Callahan, B, McMurdie, P, Rosen, M, Han, A, Johnson, A, Holmes, S. “DADA2: high-resolution sample inference from Illumina amplicon data”. Nature methods 2016; 13(7):581.

54. Katoh, K, Standley, D. “MAFFT multiple sequence alignment software version 7: improvements in performance and usability”. Molecular biology and evolution 2013; 30(4):772–780.

55. Price, M, Dehal, P, Arkin, A. “FastTree 2–approximately maximum-likelihood trees for large alignments”. PloS one 2010; 5(3): e9490.

56. Fabian Pedregosa, Gaël Varoquaux, Alexandre Gramfort, Vincent Michel, Bertrand Thirion, Olivier Grisel, Mathieu Blondel, Peter Prettenhofer, Ron Weiss, Vincent Dubourg, Jake Vanderplas, Alexandre Passos, David Cournapeau, Matthieu Brucher, Matthieu Perrot, and Édouard Duchesnay. Scikit-learn: machine learning in python. Journal of machine learning research, 12(Oct):2825–2830, 2011

57. Quast C, Pruesse E, Yilmaz P, Gerken J, Schweer T, Yarza P, Peplies J, Glöckner FO. 2013. The SILVA ribosomal RNA gene database project: improved data processing and web-based tools. Opens external link in new window Nucl. Acids Res. 41 (D1): D590–D596.

58. Abarenkov K, Henrik Nilsson R, Larsson KH, Alexander IJ, Eberhardt U, Erland S, Høiland K, Kjøller R, Larsson E, Pennanen T, Sen R, Taylor AF, Tedersoo L, Ursing BM, Vrålstad T, Liimatainen K, Peintner U, Kõljalg U. The UNITE database for molecular identification of fungi--recent updates and future perspectives. New Phytol. 2010 Apr;186(2):281–5. doi: 10.1111/j.1469-8137.2009.03160.x.

59. Martín-Hernández, R., Meana, A., Prieto, L., Martínez Salvador, A., Garrido-Bailón, E., Higes M. 2007. Outcome of Colonization of Apis mellifera by Nosema ceranae. Applied and Environmental Microbiology Oct 2007, 73 (20) 6331-6338; DOI: 10.1128/AEM.00270-0

60. Klee J, Tek Tay W, Paxton RJ. Specific and sensitive detection of Nosema bombi (Microsporidia: Nosematidae) in bumble bees (Bombus spp.; Hymenoptera: Apidae) by PCR of partial rRNA gene sequences. J Invertebr Pathol. 2006 Feb;91(2):98–104. doi: 10.1016/j.jip.2005.10.012. Epub 2005 Dec 22.

61. Yang B, Peng G, Li T, Kadowaki T. Molecular and phylogenetic characterization of honey bee viruses, Nosema microsporidia, protozoan parasites, and parasitic mites in China. Ecol Evol. 2013;3(2):298–311. doi:10.1002/ece3.464

62. Meeus I, de Graaf DC, Jans K, Smagghe G. Multiplex PCR detection of slowly-evolving trypanosomatids and neogregarines in bumblebees using broad-range primers. J Appl Microbiol. 2010 Jul;109(1):107–15. doi: 10.1111/j.1365-2672.2009.04635.x. Epub 2009 Nov 23.

